# Is the tryptophan codon of gene vif the Achilles’ heel of HIV-1?

**DOI:** 10.1101/837419

**Authors:** Élcio Leal, Fabiola Villanova, Marta Barreiros

## Abstract

To evaluate the impact of hypermutation to the HIV-1 dissemination at the population level we studied 7072 sequences HIV-1 gene vif retrieved from public databank. From this dataset 857 sequences were selected because they had associated values of CD4+ T lymphocytes counts and viral loads and they were used to assess the correlation between clinical parameters and hypermutation. We found that the frequency of stop codons at sites 5, 11 and 79 ranged from 2.8×10-4 to 4.2×10-4. On the other hand, at codons 21, 38, 70, 89 and 174 the frequency of stop codons ranged from 1.4×10-3to 2.5×10-3. We also found a correlation between clinical parameters and hypermutation where patients harboring proviruses with one or more stop codons at the tryptophan sites of the gene vif had higher CD4+ T lymphocytes counts and lower viral loads compared to the population. Our findings indicate that A3 activity potentially restrains HIV-1 replication because individuals with hypermutated proviruses tend to have lower numbers of RNA copies. However, owing to the low frequency of hypermutated sequences observed in the databank (44 out 7072), it is unlikely that A3 has a significant impact to curb HIV-1 dissemination at the population level.

---

Dear editor of PlosOne

We are pleased to submit the manuscript entitled, “*Is the tryptophan codon of gene vif the Achilles’ heel of HIV-1?*”, we present in this study results of the relationship between the hypermutation in the HIV gene *vif* and levels of CD4+ lymphocytes and levels of viral RNA.

We used 7072 proviral sequences HIV-1 gene *vif* from public databank to evaluate the impact of Apobec3 to the evolution of HIV in a population level.

First we found that tryptophan codons (TGG) of the gene *vif* targeted exclusively by RT have lower mutation rates than TGG codons target by Apobec3.

Then we used a subset of 857 sequences (sequences with information of CD+4 counts and viral loads) and found that patients harboring hypermutated viruses tend to have lower mRNA levels and higher counts of CD4+ cells

Overall our findings indicate that at individual level Apobec3-induced hypermutation a subtle effect the HIV replication. However, at the population level Apobec3 activity is minimal to restrain HIV spread because it is rare (only 0.6% of all sequence in the genbank were hypermutated).

Sincerely yours

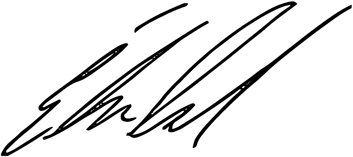

Elcio Leal

Institute of Biotechnology, Federal University of Pará;

Avenida Augusto Corrêa, 1;

CEP 66075-110;

Belém, PA, Brazil;

Phone/Fax: (55 91) 32017202 / 32017601;

**Figure#1.**
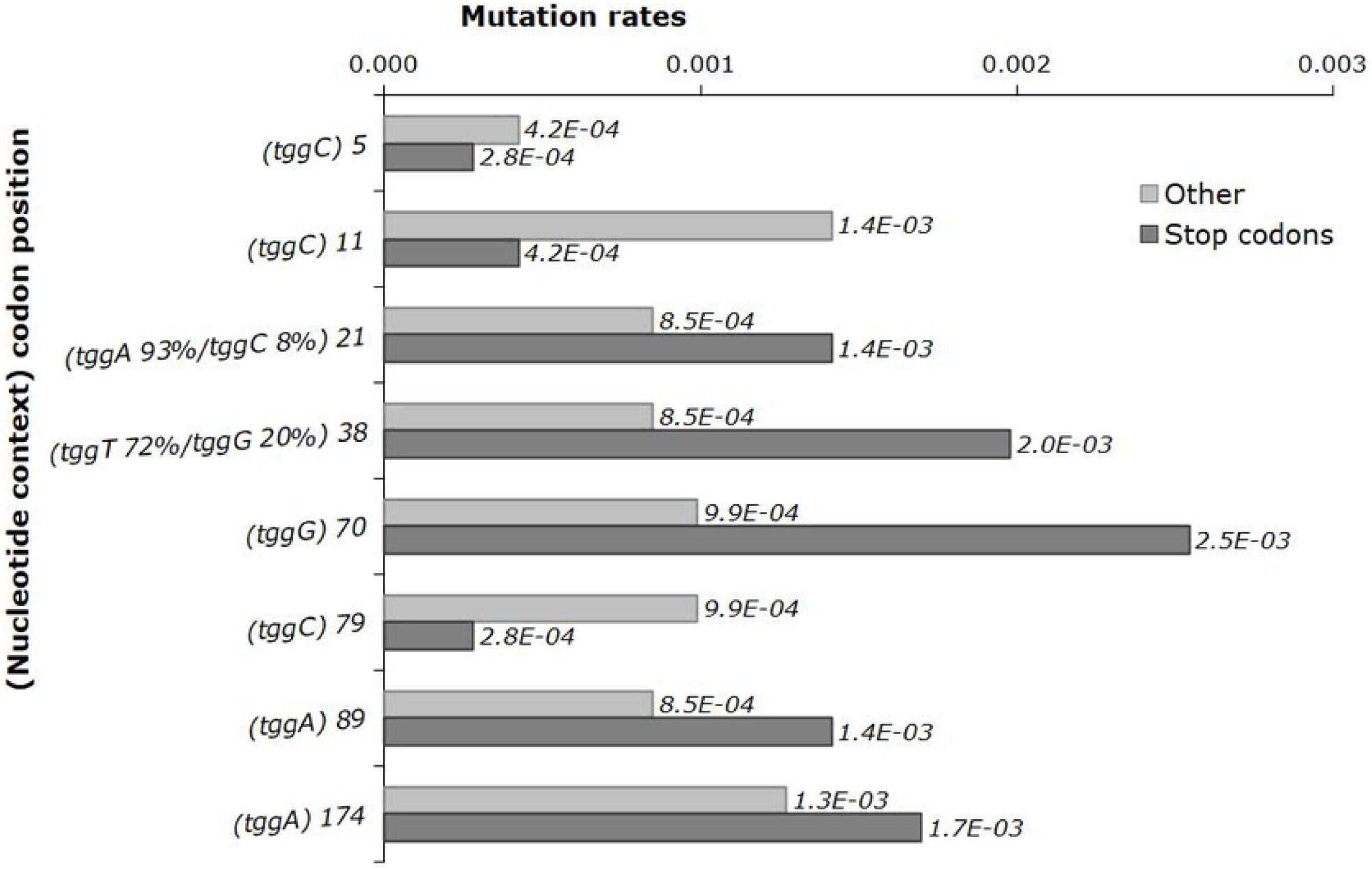

**Figure#2.**
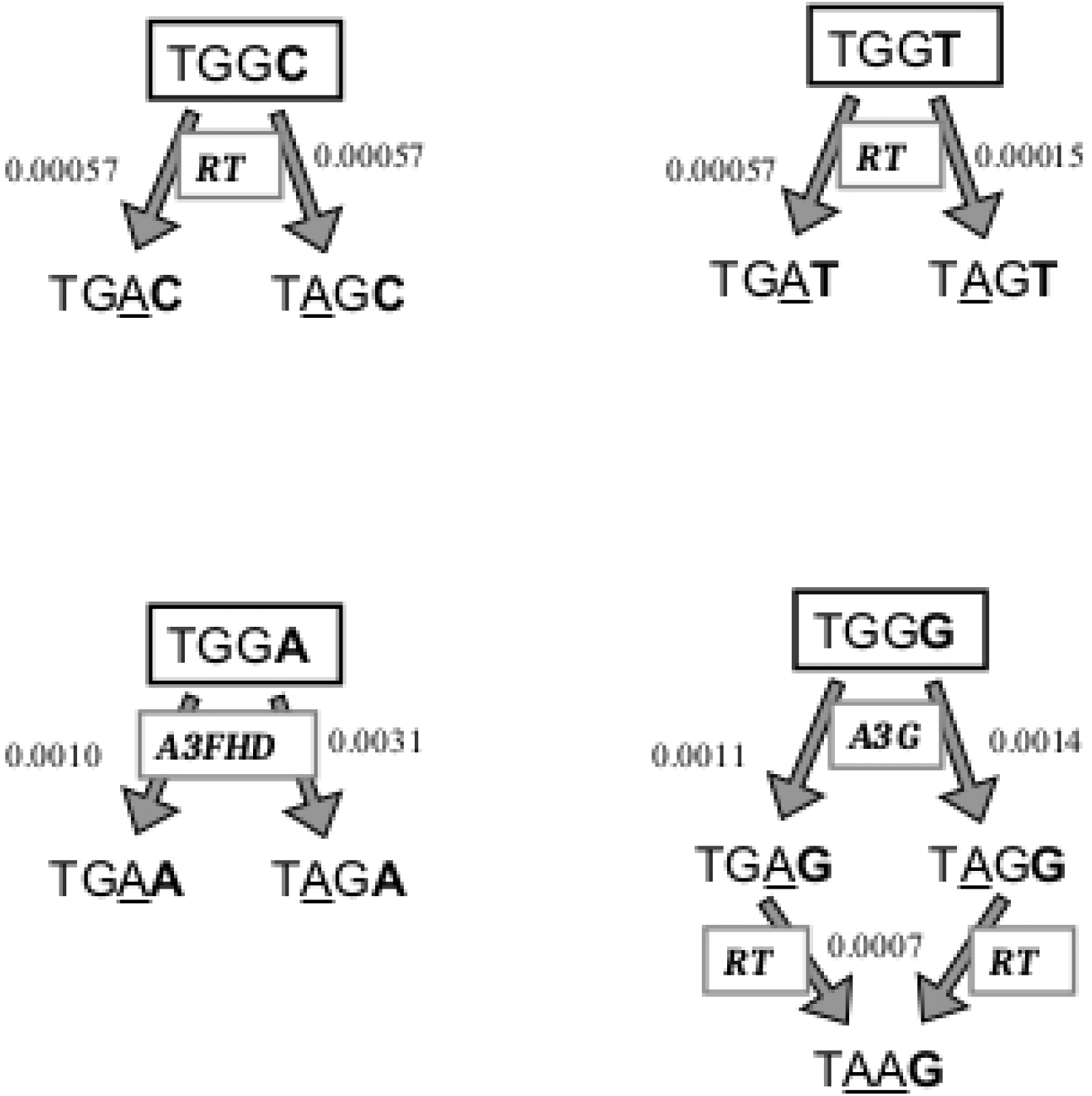

**Figure#3.**
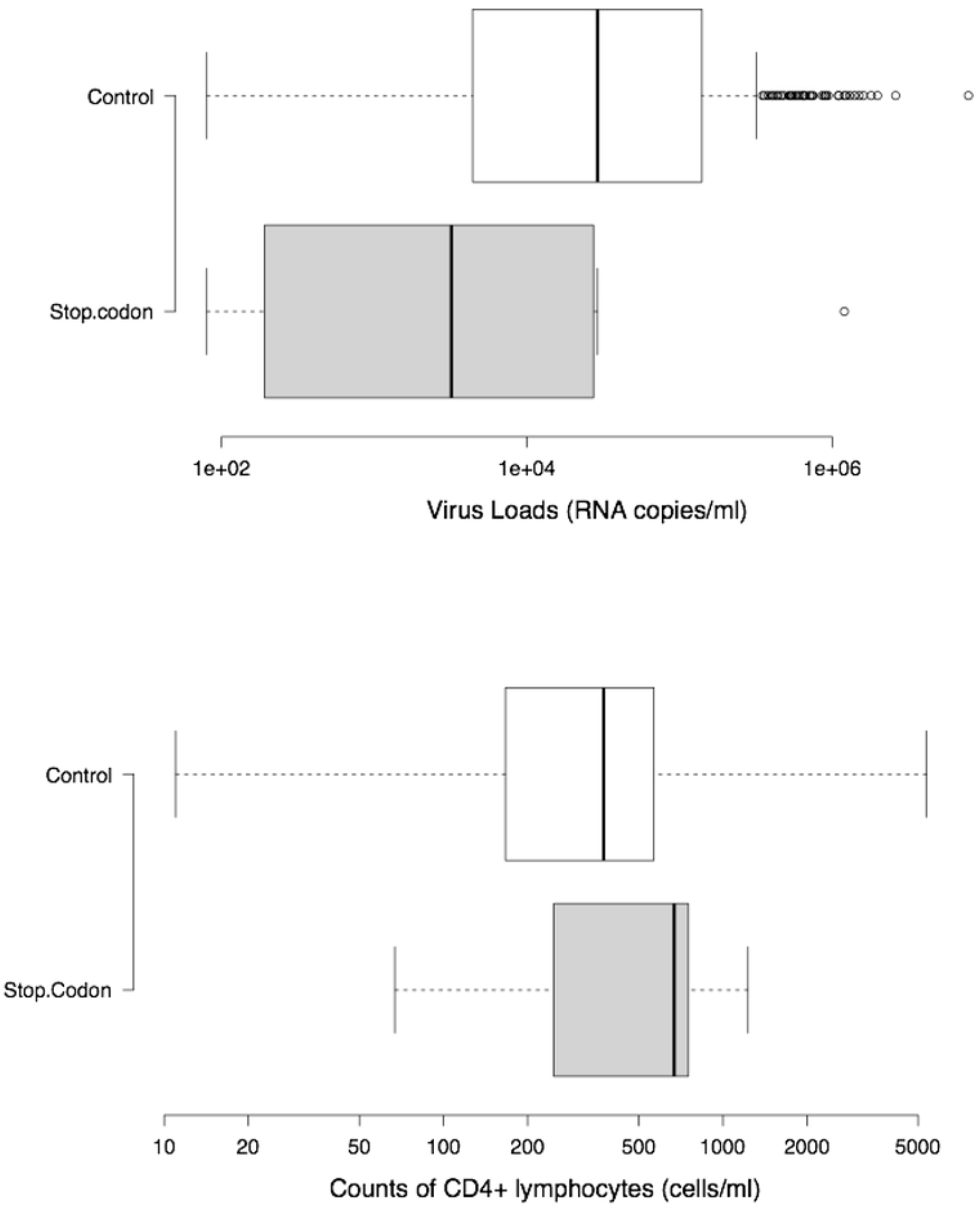

